# A comparison between Greengenes, SILVA, RDP, and NCBI reference databases in four published microbiota datasets

**DOI:** 10.1101/2023.04.12.535864

**Authors:** Camilla Ceccarani, Marco Severgnini

## Abstract

Inaccurate bacterial taxonomic assignment in 16S-based microbiota experiments could have deleterious effects on research results, as all downstream analyses heavily rely on the accurate assessment of microbial taxonomy: a bias in the choice of the reference database can deeply alter microbiota biodiversity (alpha-diversity), composition (beta-diversity), and taxa profile (bacterial relative abundances).

In this paper, we explored the influence of the reference 16S rRNA collection by performing a classification against four of the main databases used by the scientific community (i.e. Greengenes, SILVA, RDP, NCBI); the consequences of database clustering at 97% were also explored. To investigate the effects of the database choice on real and representative microbiome samples from different ecosystems, we performed a comparative analysis on four already published datasets from various sources: stools from a mouse model experiment, bovine milk, human gut microbiota stool samples, and swabs from the human vaginal environment. We took into consideration the computational time needed to perform the taxonomic classification as well.

Although values in both alpha- and beta-diversity varied a lot, sometimes even statistically, according to the dataset chosen and the eventual clustering, the final outcome of the analysis was a concordance in the capability to retrieve the original experimental group differences over the various datasets. However, in the taxonomy classification, we found several inconsistencies with taxonomies correctly assigned in only some of the four databases. The degree of concordance among the databases was related to both the complexity of the environment and its degree of completeness in the reference databases.

**IMPORTANCE:** 16S rRNA sequencing is, nowadays, the most commonly used strategy for microbiota profiling in many different ecosystems, ranging from human-associated to animal models, food matrices, and environmental samples. The ability of this kind of analysis to correctly capture differences in the microbiota composition is related to the taxonomic classification of the fragments obtained from sequencing and, thus, to the choice of the best reference database. This paper deals with four of the most popular microbial databases, which were evaluated in their ability to reproduce the experimental evidence from four already published datasets. The knowledge of the advantages and drawbacks of the database choice can be pivotal for planning future experiments in the field, making researchers aware of the repercussions of such a choice according to the different environments under scrutiny.

Moreover, this work can also shed new light upon past results, partially explaining discordant evidence.

## INTRODUCTION

Microbiota sequencing has widely spread as one of the most implemented technologies for characterizing bacterial environments. A widely used method to accomplish this characterization is to sequence amplicons targeting the hypervariable regions of the 16S rRNA gene. Due to the cost for obtaining whole metagenomes, amplicon-based sequencing is the only feasible option to screen a high number of samples at once.

Considering that most data are generated from next-generation sequencing technologies, the correct link between thousands of sequences and the established microbial taxonomic information is fundamental for metagenomic researchers. Hence, it is crucial to use correct and reliable reference databases, as incorrect assignments can be deleterious to all downstream analyses.

The influence of the taxonomy database on the bacterial classification has been explored using different types of mock communities (1,2), reference type strains (3), stool samples from healthy volunteers (2) or directly comparing taxonomies within each database (4); compared databases include Greengenes, SILVA (1-4), RDP (2,4), EzBioCloud (1,3), NCBI, OTT (4), the genomic-based 16S rRNA Database GRD, and The All-Species Living Tree (LTP) (2). All of them stressed the importance of the correct choice of the reference database, as well as their advantages and disadvantages. However, none but one of them (2) employed real samples or already published datasets.

In this study, we aimed at not only unraveling the differences between the most used databases (i.e.: Greengenes, SILVA, RDP, NCBI) on real datasets but also help discern the implementation of a complete database rather than a clustered one (Greengenes and SILVA at 97% *vs* their 100% identity files version). It is true that several online tools and user-friendly software, like Galaxy, MEGAN and QIIME2 (5-7), have been crafted to provide even more simplicity for any researcher struggling to analyze its sequencing reads, but choosing among the variety of microbial databases for classifying bacterial species could still be puzzling even for an expert eye.

Although it has not been updated since 2013, the Greengenes 16S rRNA gene database (8) is one of the first and most commonly used databases, even in the newer studies, both for OTU and ASV/zOTU workflows, thanks to the inclusion in popular metagenomic analyses packages such as QIIME and QIIME2 (5,9). It is a collection of the 16S rRNA gene records in the GenBank database (10) with descriptive fields and taxonomic assignment, a chimera screening, and ARB (11) compatibility features. Contrariwise, the SILVA database (12) has become widely used as it is constantly updated, and it is one of the default databases in the QIIME2 package. Through a manual curation approach based on the phylogenetic tree, SILVA has its taxonomy annotations updated for Bacteria, Archaea, and Eukarya species. The Ribosomal Database Project (RDP, 13) contains aligned and annotated bacterial and archaeal small subunit rRNA gene sequences, the majority of which are not complete over the whole 16S gene. Most of these are derived from the International Nucleotide Sequence Database Collaboration (INSDC) databases and sequencing PCR amplification products, whereas a small number of older entries derive from isolated rRNA reverse-transcriptase sequencing. Roughly 85% of bacterial sequences in RDP are from directly-isolated DNA from environmental samples, and only a small percentage originate from cultured organisms. The NCBI Taxonomy database (14) is known as the standard nomenclature and classification repository for the INSDC, incorporating databases like GenBank, ENA (EMBL), and DDBJ (15). Like SILVA, this database is manually curated by a group of scientists exploiting the most recent taxonomic literature to confirm or update phylogenetic taxonomy.

In order to provide the research community with the clearest possible evaluation, we performed a classification analysis with these databases on four datasets collected from both animal and human subjects, and from disparate districts. The variety of the bacterial environments used in this study allowed us to test those databases with more depth than the comfort zone given by the widely used and characterized gut microbiota.

## MATERIALS AND METHODS

### Dataset selection

The involved studies were selected according to the high differences between the cohorts involved and according to the different nature of the samples sequenced. In order to obtain an extensive observation and to provide information useful for a large audience of researchers, we chose four already published studies from our research group, all with publicly available sequence data, that allow this project to range from murine models to livestock to human beings, and from the most used types of samples (i.e. stool) to the less common (e.g.: milk).

Animal datasets:

- The murine model (“mouse infection”) was the one described in Majlessi et al (16). A comparative study of the gut microbiota composition in B10.D2 mice, pathogen-free (n=6) or naturally colonized with *Helicobacter hepaticus* (n=6) was performed.
- The bovine milk (“milk microbiota”) dataset was the one analyzed by Cremonesi et. al. (17) and compared the microbiota composition of milk quarter samples between two cow breeds: the highly productive Holstein Friesian (HF, n=20), and the autochthonous Italian breed Rendena (REN, n=10). Among the original 117 milk samples, only those collected at time point T4 were chosen as representative.

Human datasets:

- Stool samples from a study comparing gut microbiota composition of people at different stages of life (“extreme longevity”), performed by Biagi et. al. (18). We selected the young-adults (age: 22-48y, n=15) and the semi-supercentenarians cohorts (age: 105-109y, n=24), as a representation of a clear distinction of microbiota patterns based on age.
- Vaginal swabs (“vaginal microbiota”) from reproductive-aged women affected by different vulvo-vaginal infections, studied by Ceccarani et. al. (19). Of the four cohorts involved, for this present study, we selected the healthy controls (HC, n=21) and women with bacterial vaginosis (BV, n=20).

### Data availability

Three out of four datasets are available in the SRA database (https://ncbi.nlm.nih.gov/sra) with the following accession numbers: PRJNA523312 (vaginal microbiota), PRJNA414712 (milk microbiota), PRJNA320730 (mouse infection). The “extreme longevity” dataset is available on MG-Rast (https://www.mg-rast.org/index.html) with project ID mgp17761.

### Microbiota profiling and Bioinformatic analysis

The data analysis procedures were the same as those employed in the original publication, except for the use of different reference databases for the taxonomy assignment and the phylogenetic tree building. The 16S rRNA raw sequences obtained in each project were merged using Pandaseq (release 2.5; 20), filtered by trimming stretches of 3 or more low-quality bases (quality < 3), and discarding the sequences whenever they were shorter than 75% of the length of the original fragment. As in the original studies, all bioinformatic analyses were conducted using the QIIME pipeline (release 1.9.0; 21), clustering filtered reads into Operational Taxonomic Unit (OTUs) at 97% identity level and discarding singletons as possible chimeras. Taxonomic assignment was performed via the RDP classifier (22), using 0.5 as the confidence threshold, against each of the following databases:

- Greengenes (100%), release 13.5; Greengenes 97%, release 13.8 (ftp://greengenes.microbio.me/greengenes_release/)
- SILVA (100%), and SILVA 97%, both release 132 (https://www.arb-silva.de/no_cache/download/archive/release_132/)
- RDP, release 11.5 (https://rdp.cme.msu.edu)
- NCBI, release June 17th, 2018 (ftp://ftp.ncbi.nlm.nih.gov/blast/db/)

The four databases contain a greatly different number of sequences, ranging from 19’890 (NCBI, release 2018), to 1’262’986 (Greengenes), to 2’090’668 (SILVA 132), and to 3’196’041 (RDP).

Alpha-diversity was computed through the QIIME pipeline using the Chao1, the number of OTUs (“observed species” index), Shannon diversity, and Faith’s Phylogenetic Diversity whole tree (PD whole tree) metrics. To compare the microbial community structure of the subjects for the beta-diversity analysis, weighted and unweighted UniFrac distances (23) were used. For creating the OTUs phylogenetic tree, for each database, the specific set of multiply-aligned sequences was used as a template for the PyNAST alignment method. Full details on the procedures used for preparing the databases for the analyses are provided as Supplementary Methods.

All statistical evaluations and visualizations were performed in Matlab (v. 2008, Natick, MA, USA). Statistical significance of the alpha-diversity differences was assessed by non-parametric Monte Carlo-based tests, whereas beta-diversity differences were determined through a permutation test with pseudo-F-ratios (“adonis” function from R package “vegan”, version 2.0-10. Pairwise relative abundance analysis was performed using a non-parametric Mann–Whitney U test, considering a p-value <0.05 as the threshold for significance.

## RESULTS

### Computational performances

Analyses have been performed on an HP ProLiant DL580 server with 4 × 14 CPUs at

2.0 GHz and 1TB of RAM (Table 1).

**Table 1.**
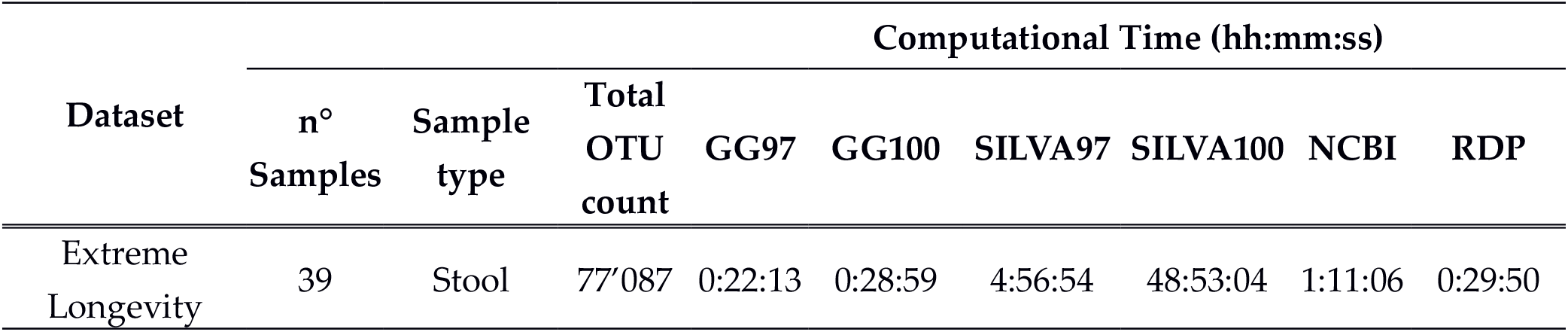
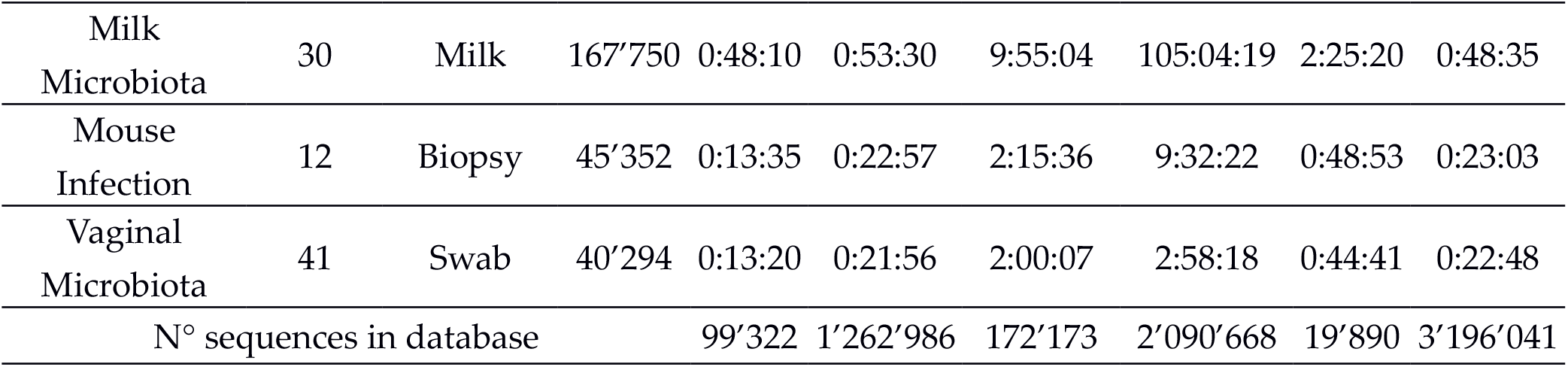
Computational time required by our server for performing the taxonomic classification with the RDP classifier for the four databases over the four datasets. Where available, performances on 97% clustered and raw (100%) taxonomies are reported as well.

Greengenes complete (100%) database performed 1.10-1.69 times slower than the 97% version in each dataset analyzed, whereas SILVA 100% identity database performed the taxonomy assignment step even slower (9.88x, 10.59x, 4.24x, and 1.48x for extreme longevity, milk microbiota, mouse infection and vaginal microbiota, respectively).

Time necessities were also different between Greengenes, SILVA, RDP, and NCBI databases. On the average of all computational times normalized for the total OTU count, RDP was the fastest, 1.01 times faster than Greengenes, 0.02x than SILVA, and 0.44x than NCBI. Greengenes and the NCBI databases had comparable computational times, while SILVA was the slowest one: 53.09x *vs* Greengenes; 23.27x *vs* NCBI; 52.85x *vs* RDP. Lastly, the average process duration (minutes per OTU) was, from the fastest to the slowest database, 4.36×10^−4^ for Greengenes, 4.38×10^−4^ for RDP, 9.96×10^−4^ for NCBI, 2.32×10^−2^ for SILVA.

### SILVA and Greengenes databases at different identity

First, we sought to highlight any eventual difference in the microbiota assessment deriving from the taxonomic classification based on the databases (SILVA and Greengenes) with taxonomy files grouped on identity percentage: 97% and 100%.

### Alpha-diversity evaluation

As expected, biodiversity evaluated with non-phylogenetic-based metrics did not change. Contrariwise, Faith’s phylogenetic diversity (PD_whole_tree) values slightly differed: for the SILVA database, 97% identity values were higher than those at 100%, whereas for the Greengenes database we observed the opposite. When comparing the two databases versions, two datasets (mouse infection and extreme longevity) had significantly higher Greengenes estimates than their SILVA counterpart (p<0.002, unpaired t-test); milk microbiota showed a non-significant increased biodiversity tendency in Greengenes compared to SILVA; vaginal microbiota reported a 97% SILVA biodiversity significantly higher (p<0.0001) than both Greengenes-based and 100% SILVA estimates (Supplementary Table 1). However, independently from the identity percentage of the database employed, significant differences in alpha-diversity values over experimental categories were retained (Supplementary Figure 1).

### Beta-diversity evaluation

Microbiota composition through beta-diversity analysis showed that the database choice (SILVA or Greengenes) and the different identity percentage (97% or 100%) did not change the significant distance between microbial profiles originally found for all four published datasets, both for unweighted and weighted Unifrac distances (p-value <0.004, adonis test) (Figure 1). The 97% or 100% databases choice had a clear effect on the weighted Unifrac distance distribution among experimental group: Greengenes 97% and 100% distributions resulted statistically different (p<0.001) among all but mouse infection dataset, whereas SILVA 97% and 100% distributions were significantly different (p<0.001) among all four datasets. Notably, these differences were particularly evident in the mouse infection dataset (average distances of 0.17 and 0.30 for SILVA 100% and 97%, respectively) and in the vaginal microbiota dataset (0.22 and 0.52 for SILVA 100% and 97%), with the 100% database resulting in more similar profiles. Unweighted Unifrac distances among samples resulted unchanged, except for the extreme longevity and the vaginal microbiota datasets (Supplementary Table 2).

**Figure 1.**
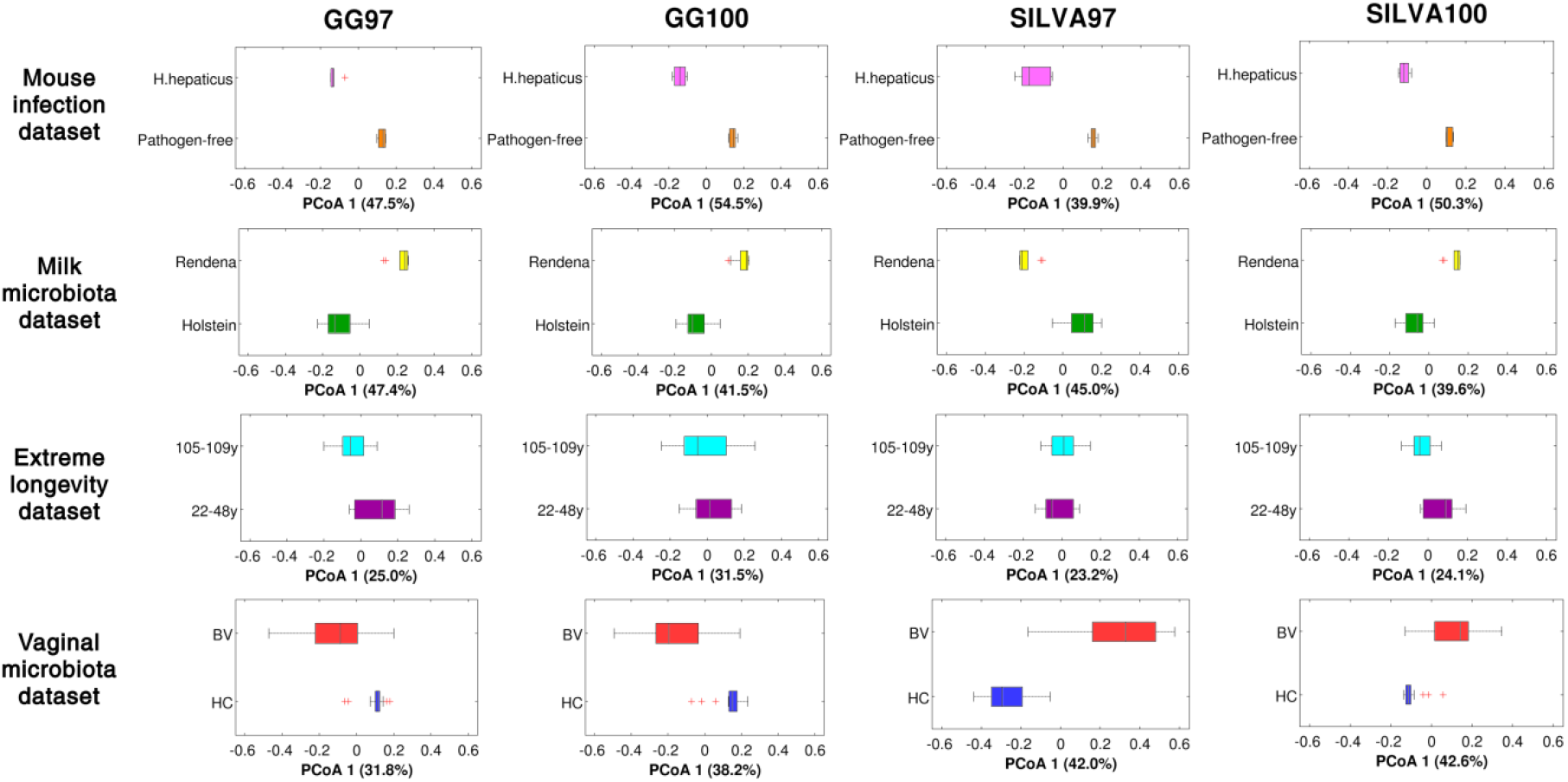
Beta diversities for the raw and clustered reference databases. Horizontal boxplots of the distribution of the Principal Coordinate Analysis (PCoA) on weighted Unifrac distances, using SILVA and Greengenes (“GG”) databases at 97% or 100% for the four datasets considered. For all the datasets, the first principal coordinate (PCoA 1) was associated with the experimental class. Percent contribution of the coordinate to the total variance is reported below each graph. For each boxplot the median is reported.

### Bacterial taxonomy characterization

Relative abundances at genus level were generally akin when comparing 97% to 100% identity databases, with an average difference over the main taxa of <0.2% in both Greengenes and SILVA over all the four datasets.

The mouse infection dataset was characterized by the highest disagreement, mainly due to the unclassified bacterial taxa: *Other Clostridiales* (+1.4% on average), *Unclassified Clostridiales* (−6.5%), *Unclassified Lachnospiraceae* (+4.6%) for the Greengenes database; *uncultured bacterium* (*Muribaculaceae* family) (+9.4%) and *Unidentified muribaculaceae* (−7.9%) for the SILVA database.

In the milk microbiota dataset, the 97% database reflected in a slight overestimation of the genus *Streptococcus* (<1% in both Greengenes and SILVA), whereas *Lactobacillus* was underestimated (−0.3%) in Greengenes and overestimated (+1.3%) in SILVA.

For the extreme longevity dataset, several groups presented differences, such as *Unclassified Ruminococcaceae* (−4.8%), *Unclassified Lachnospiraceae* (−1.2%), *Ruminococcaceae (other)* (+2.0%) for the Greengenes database; the SILVA database reported an even lower range of disagreement between 97% and 100% versions (maximum difference: *Lachnoclostridium*, -0.1%).

In the vaginal microbiota dataset, Greengenes 97% database reported an increased abundance of *Unclassified Ruminococcaceae* (+2.3%) and a decreased abundance of *Faecalibacterium* (−1.7%); the SILVA database, instead, had less significant variations (all <0.3%) (Figure 2).

**Figure 2.**
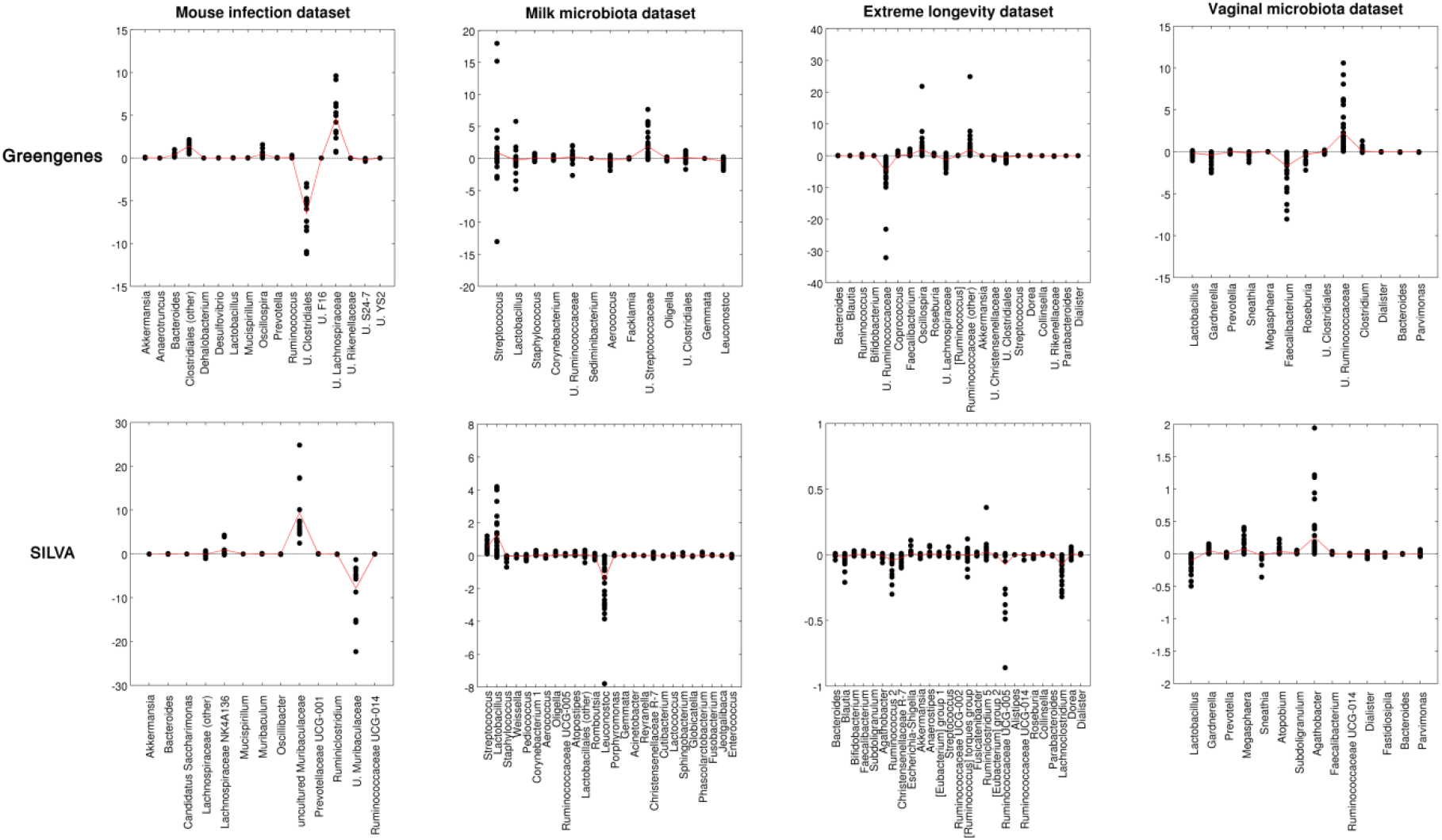
Relative abundance comparison for the raw and clustered databases. Dot plots of relative abundance differences between databases. Each subplot represents a combination of dataset and database. For each genus, the difference between relative abundances for 97% and 100% similarity databases is plotted as a black dot for each sample (n=12, 30, 39 and 42 for mouse infection, milk microbiota, extreme longevity, and vaginal microbiota datasets, respectively). Positive differences indicate a higher abundance in the 97% database and negative differences the opposite. The average difference is represented as a solid red line.

### Comparison between SILVA, Greengenes, RDP, and NCBI databases

We evaluated microbiota composition differences deriving from the different taxonomic classifications obtained with the four major databases considered for our study: Greengenes, SILVA, RDP, NCBI. Since RDP and NCBI do not have a 97% clustered version, 100% SILVA and Greengenes versions were used.

### Alpha-diversity evaluation

For all four datasets, we noticed that biodiversity estimates were highly variable when comparing the PD_whole_tree values. At least one database resulted in significantly different estimation values (p≤ 0.002, one-way ANOVA). Some differences were notable: mouse infection and extreme longevity datasets reported significantly different pairwise comparisons, as the mean estimation of each database was different from all the others (p≤ 0.045, Tukey HSD test). SILVA was characterized by the lowest biodiversity in all databases, whereas RDP (mouse infection and vaginal microbiota datasets) and NCBI (milk microbiota and extreme longevity dataset) reported the highest ones. (Figure 3)

**Figure 3.**
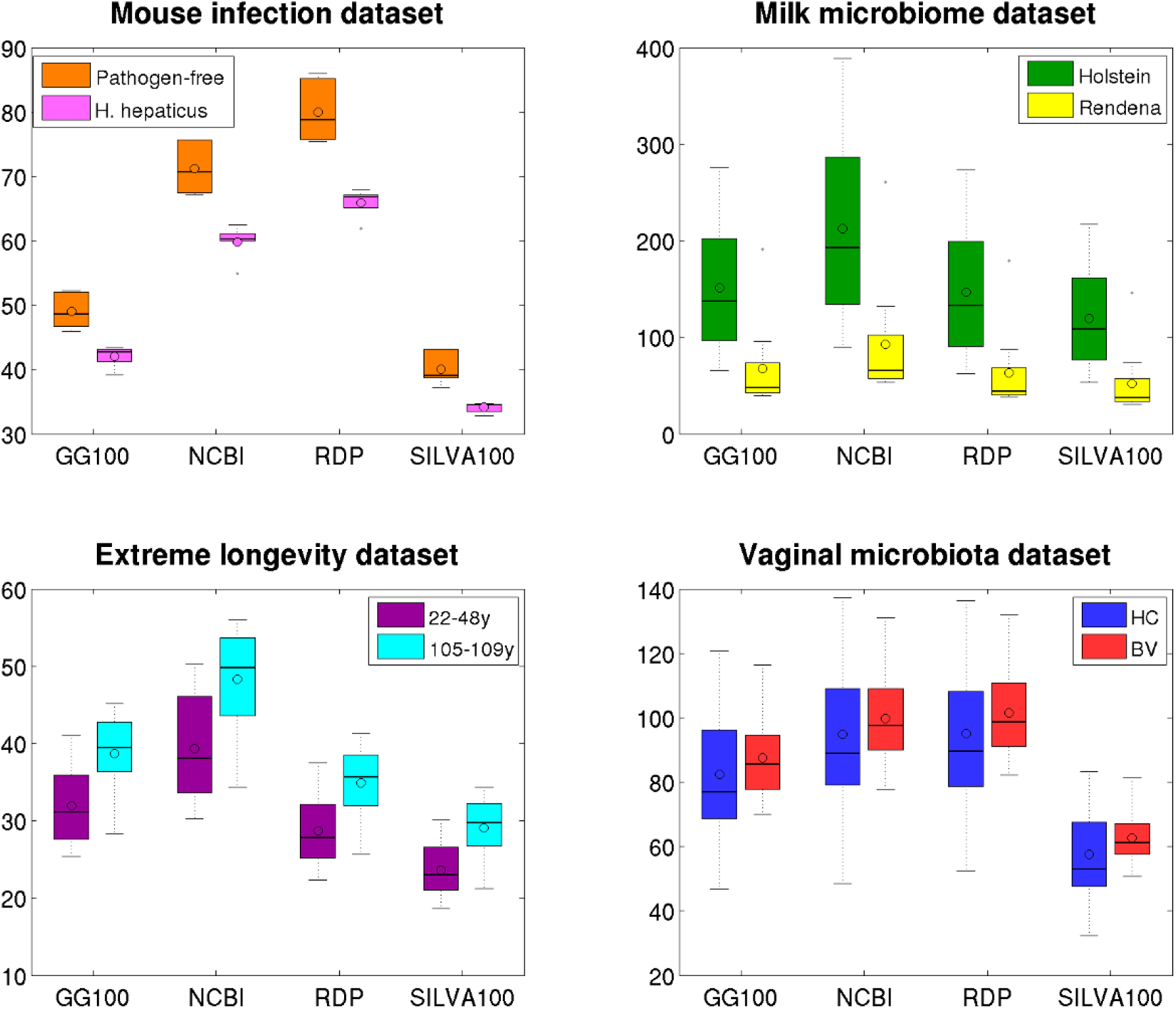
Alpha-diversity estimates for the four reference databases. Boxplots of alpha-diversity estimations according to Faith’s phylogenetic diversity metric (PD whole tree) for the four datasets analyzed and the four reference databases. In each boxplot, the median value is represented as a solid black line, whereas the mean is represented with a circle.

### Beta-diversity evaluation

While retaining the differences described in the original papers, our beta-diversity analysis reported different microbial profiles for all experimental categories in the datasets, for both unweighted and weighted Unifrac distances (p-value <0.004, adonis test) (Figure 4).

**Figure 4.**
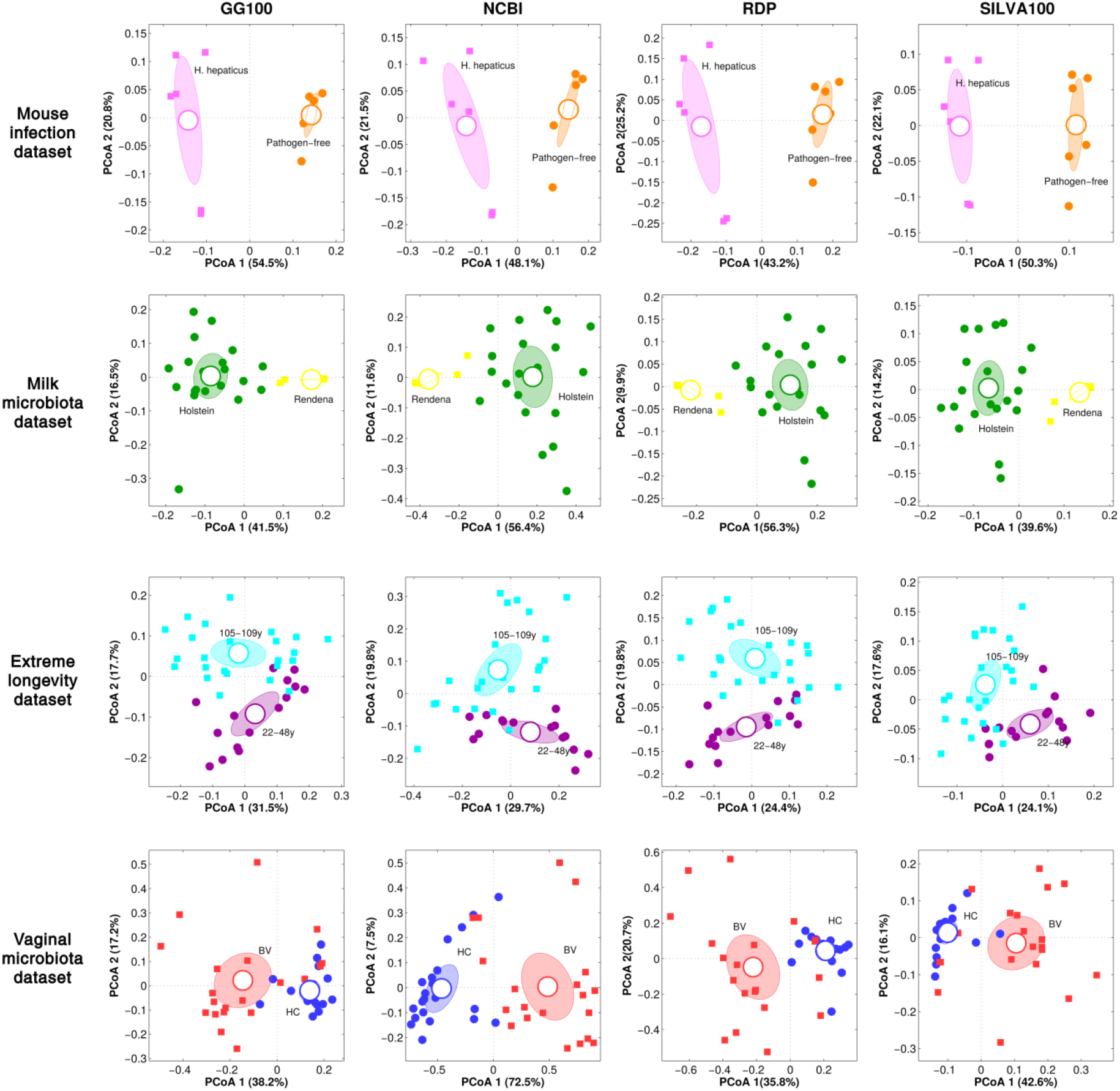
Beta-diversity evaluation for the four reference databases. Principal Coordinate Analysis (PCoA) plots based on weighted Unifrac distances among samples for each of the four datasets and databases analyzed. The first principal coordinate (PCoA 1) was reported as associated with the experimental class. Each point represents a sample; ellipses are the 95% SEM-based confidence intervals; elements are colored according to the experimental condition.

SILVA-derived distances were found consistently lower (p<0.001, Tukey HSD test) than in the other databases, particularly evident in the vaginal microbiota dataset (average distance: 0.22, 0.35, 0.57, and 0.50 for SILVA, Greengenes, NCBI, and RDP, respectively); contrariwise, NCBI (extreme longevity, milk, and vaginal microbiota datasets) and RDP (mouse infection dataset) databases were the ones with higher distances. While this behavior was typical of the weighted distances, differences in unweighted Unifrac were only seen with the extreme longevity and vaginal microbiota datasets (Supplementary Table 3).

### Bacterial taxonomy characterization

To understand whether and how the choice of a database may influence the taxa observed in the original papers, we compared the relative abundance according to the four databases analyses for all the datasets. We reported and discussed genus-level findings for easier comparisons with the study of origin main results.

In the mouse infection dataset, all databases reported a significant (p<0.05) increase of the genera *Bacteroides* and *Helicobacter*, and a significant decrease of *Prevotella* in *H. hepaticus*-infected mice; *Akkermansia* (p>0.05) was consistently detected by all databases. Many other bacteria, however, displayed a non-consistent classification, particularly involving *Muribaculum* (nearly exclusively in NCBI), *Unclassified Muribaculaceae* (SILVA), and a further group classified as *Unclassified S24-7 / Uncultured bacterium (Muribaculaceae) / Unclassified Porphyromonadaceae* present in all databases but NCBI. Classifications with RDP and Greengenes also reported a decrease in unidentified members of *Clostridiales*, although at very different abundances (>20% and <5%, respectively); Greengenes and SILVA were concordant in reporting the decrease of members of the *Ruminococcaceae* family, whereas NCBI only indicated a *Barnesiella* increase. (Figure 5A) Characterization of the milk microbiota dataset was highly consistent among all the four databases: they all reported significantly increased abundances of the genera *Streptococcus, Lactobacillus, Pediococcus, Leuconostoc*, and *Enterococcus* in Rendena cows, and decreased abundances of *Bradyrhizobium/Nitrobacter, Staphylococcus, Aerococcus, Corynebacterium*, unclassified *Ruminococcaceae*. Average relative abundances were highly similar among all databases, accounting together for 83.4% - 91.5% (depending on the database) and for 48.5% - 51.2% of the average relative abundance in Rendena and Holstein breed, respectively. Moreover, among increased bacteria in the Holstein breed, *Facklamia, Oligella*, and unclassified *Clostridiales* were detected by all but the SILVA database, whereas *Weissella, Romboutsia, Atopostipes, Porphyromonas*, and *Acinetobacter* were concordant on all databases but Greengenes. Overall, for this dataset, when two or more datasets congruently classified a taxon, relative abundances were similar. (Figure 5B) In the extreme longevity dataset, *Bacteroides, Faecalibacterium, Escherichia-Shigella, Akkermansia, Eggerthella*, and *Anaerotruncus* were consistently classified by all databases, with the same increasing and decreasing trends of the two experimental groups; contrariwise, *Blautia, Dialister*, and *Anaerostipes* were commonly classified by three resources. Only NCBI and RDP detected *Dorea;* only NCBI and SILVA *Fusicatenibacter*.

**Figure 5.**
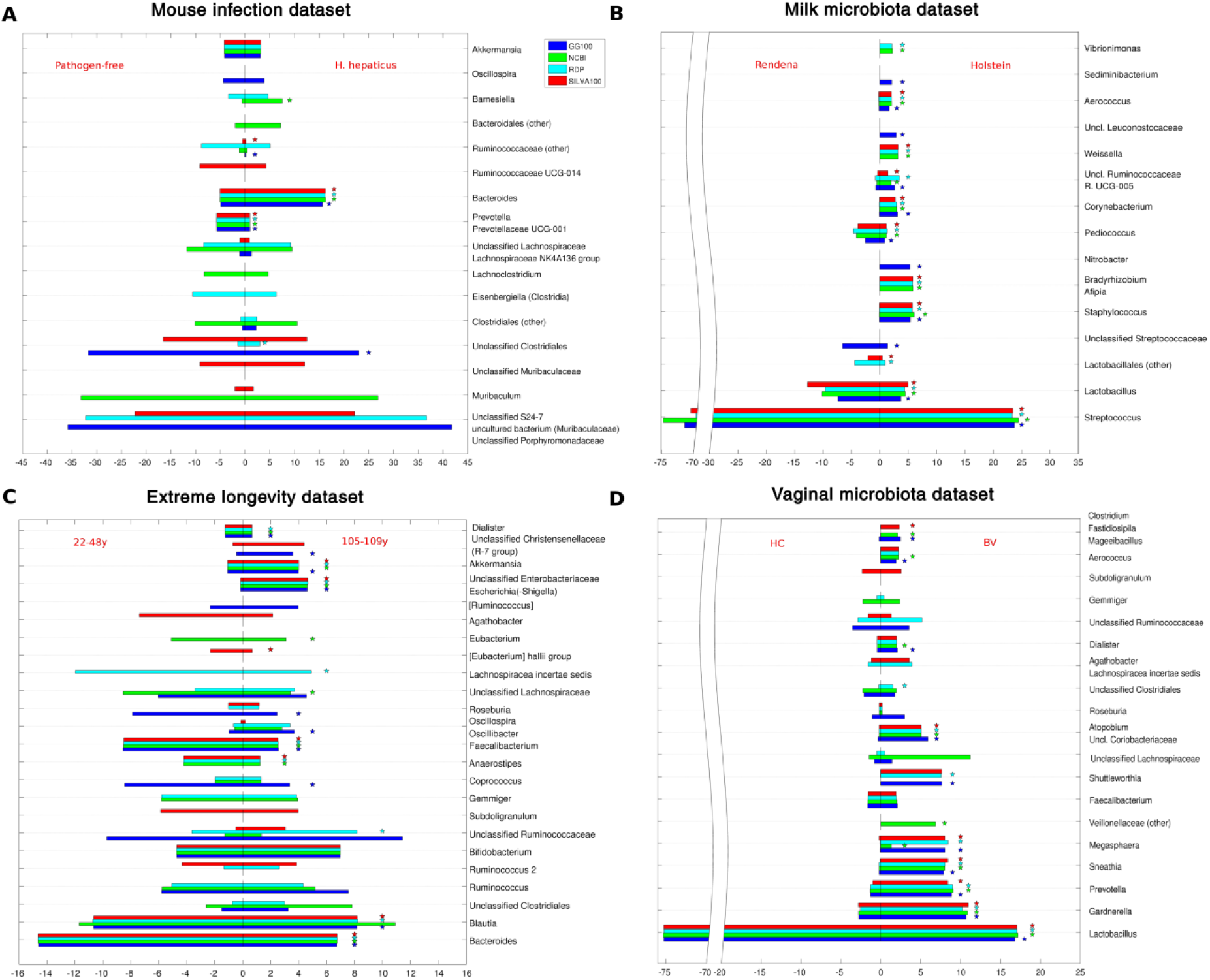
Relative abundance comparison for the four reference databases. Barplots of average relative abundances at genus level for the four datasets. For each genus, the average abundance for each specific database is reported as differently colored bars. In order to represent both experimental conditions, each plot was divided in two halves. Statistically significant differences are evidenced by a star above the rightmost bars.

Other exclusively detected genera, all increased in the 105-109y group: *Oscillospira* and members of the *Christensenellaceae (*by Greengenes); *Ruminococcaceae* families (by NCBI). Similarly, three taxa (*Coprococcus, Roseburia, Unclassified Lachnospiraceae*), all increased in the 22-48y group, were found only by Greengenes (the former two) and NCBI (the latter). Consequently, their relative abundances were also different according to the database used: e.g. *Roseburia* in the 22-48y group accounted for about 8% in Greengenes, and <1% in the others; *Coprococcus* in the 22-48y group was 8% in Greengenes and <2% in the others. (Figure 5C)

Lastly, in the vaginal microbiota dataset, all the four databases consistently classified the main bacterial genera and their changes, such as the statistically significant *Lactobacillus* (depleted in BV-affected women), *Gardnerella, Prevotella, Sneathia, Megasphaera, Atopobium* (increased in BV), plus *Faecalibacterium* (unchanged). These bacterial relative abundances were nearly identical over the databases, except for *Megasphaera* that NCBI accounted for only 1.3% in BV women (as compared to about 8% in all remaining databases). Among the less abundant genera, *Dialister* and *Aerococcus* were both consistently classified by all databases, whereas *Shuttleworthia* (average rel. ab. 7% in BV *vs* <0.1% in HC) was nearly absent in NCBI. Finally, *Roseburia* (2.0% in Greengenes *vs* 0.2% with the other three databases), *Gemmiger* (2.3% in NCBI *vs* 0.2% with the others), and *Subdoligranulum* (2.4% in SILVA *vs* <0.1% with the others) had fluctuating abundances across databases. (Figure 5D)

### Concordance among the four databases

In order to highlight concordant/discordant classifications, we compared OTUs classification of each database against the others (Figure 6, Supplementary Figure 2).

**Figure 6.**
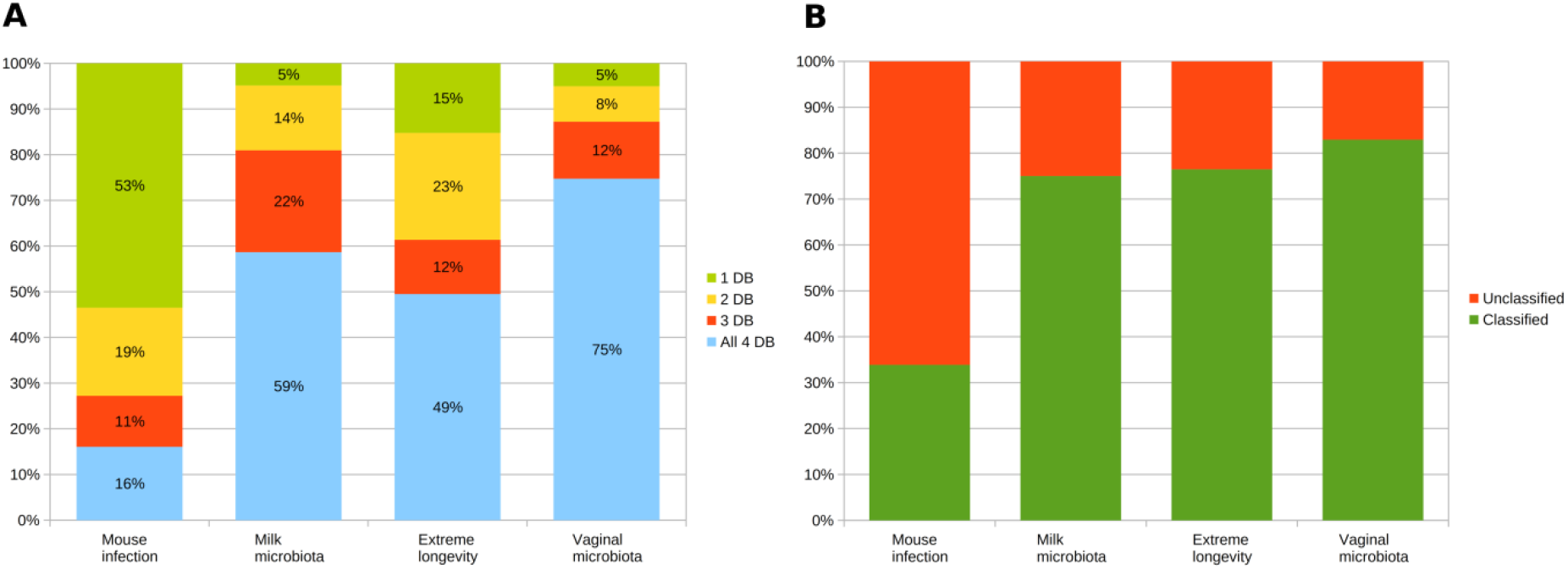
Classification concordance among the four databases. (A) Stacked barplots representing the relative abundances (genus level) that were classified all discordantly (“1 DB”) or concordantly by 2 or more databases (“2 DB”, “3 DB”, “4 DB”) for each of the four datasets; (B) Barplots representing the proportion of relative abundance related to taxa classified or unclassified at genus level. “Unclassified” means unclassified at genus level in at least one of the taxonomy databases.

The mouse infection dataset reported the lowest concordance among databases: 53.5% of the average relative abundance was made up of classifications different for all databases, with 52.4% linked to unclassified genus-level taxa. 24.4% of the average total relative abundance belonged to a group classified differently for each database: *Unclassified Porphyromonadaceae* (RDP), *Muribaculum* (NCBI), *Uncultured bacterium* (SILVA), *Unclassified S24-7* (Greengenes). Contrariwise, 19.3% of the average relative abundance was related to taxa identically classified by two databases, 11.2% was related to taxa concordant over three databases and the remaining 16.1% related to OTUs identically classified by all databases: among these, *Bacteroides* accounted for the vast majority (10.3%), followed by *Mucispirillum* (0.8%) and by *Lactobacillus* (0.4%).

Evidence from the milk microbiota dataset provided that 58.7% of the relative abundance was identically identified by all databases, deriving mainly from *Streptococcus* (37.1%), *Lactobacillus* (4.7%), and *Staphylococcus* (3.5%); a further 22.3% was related to genera identically classified by three databases (15.9% in all but Greengenes), whereas 14.2% was congruently classified by two databases. Only 4.9% of the total relative abundance was of bacteria classified differently in all databases, 4.6% of which was due to “unclassified” genus-level taxa.

In the extreme longevity dataset, 49.5% of the relative abundance was identically classified by all databases, including many main gut microbiota members, such as *Bacteroides* (9.6% on average), *Bifidobacterium* (6.1%), *Blautia* (9.0%), *Faecalibacterium* (4.8%), *Akkermansia* (2.8%), and *Roseburia* (0.6%). A further 11.9% was due to taxa concordantly classified by three databases (9.3% identically identified in all but Greengenes): this group was characterized by some members of the *Blautia, Roseburia*, and *Bacteroides* genera. Concordance between the two databases summed up to 23.4% of relative abundance, whereas, in this dataset, the number of differently classified taxa (15.2%) was significantly lower than that of the mouse infection dataset, with no clear-cut indication of a systematic problem of misclassification, since the most abundant taxon accounted for only 2.5%.

Finally, within the vaginal microbiota dataset, all databases were concordant in the main genera classification: *Lactobacillus, Gardnerella, Prevotella, Sneathia, Megasphaera* (up to 74.7% of the total relative abundance). Concordance for three databases was recorded for 12.5% of the relative abundance, involving *Atopobium* (2.5%), *Megasphaera* (3.3%), and *Shuttleworthia* (3.6%). Moreover, 7.8% of the relative abundance was due to taxa identically classified by two databases, while 5.0% was differently classified in all.

Supplementary File 1 reports all the OTUs and the associated taxonomies in the four databases for each dataset, together with the full summary tables.

## DISCUSSION

Choosing the reference database to classify OTUs against, in microbiota profiling analyses, usually follows more practical than theoretical reasons: admittedly, due to the different size and number of reference sequences, it has direct consequences on the classification duration time. The database choice is often dictated by the pipeline used: for example, the QIIME pipelines suggest only Greengenes or SILVA, mothur (24), allows either Greengeens, SILVA, or RDP, and MEGAN (6) can use both NCBI or SILVA.

Determining the reference database can profoundly influence all resulting analyses: bacteria will be clustered differently, with the consequent risk of non-consistent results across databases. The identity grouping percentage plays an important role as well, as both Greengenes and SILVA provide different identity levels, with 97% being the most popular: strains belonging to the same bacterial species are believed to have <3% difference in their 16S rRNA sequence (25), an aspect that has been recently debated (26).

The database choice impacts not only on the taxonomic composition (especially at genus level) but on all phylogenetic-based evaluations too, both on alpha-diversity metrics and beta-diversity distances, since each database has its specific multiple-aligned reference set as a guide for assembling the phylogenetic tree.

We here explored the influence of the choice of both the identity percentage and the reference database on four already published datasets, each referring to a different microbiota source: mouse and human stools; human vaginal mucosa; and bovine milk.

### Diversity estimates and relative abundances

Alpha- and beta-diversity analyses with all databases resembled the differences described in the original papers, and this applies to the percentage of identity as well. It’s worth noting that, despite this concordance in the capability to retrieve the original differences and the robustness over datasets, values in both analyses varied a lot, sometimes even statistically, according to the source. This result suggests that differences among the four databases are indeed present and that are replicated over the samples, without significantly changing the ultimate evidence about the experimental categories.

However, when considering the taxonomic profiles, a more complex picture was evidenced, with 97% and 100% databases performing nearly the same in all the datasets, with the only differences to be accounted for genus-level unclassified taxa.

### Comparison with the original papers

The original paper by Majlessi and colleagues (16) only reported family-level differences: *Bacteroidaceae, Helicobacteraceae*, and other unclassified members of *Bacteroidales* enriched in *H. hepaticus*-colonized mice; *Unclassified Clostridiales, Ruminococcaceae, Lachnospiraceae*, and *Prevotellaceae* reduced. These results were substantially confirmed in our analysis, while all databases congruently reported genus-level alterations in *Prevotella, Helicobacter, Bacteroides. Unclassified Bacteroidales* was reported only by two databases, due to the different classification of the *Muribaclulum* genus, similarly to the reported decrease in unclassified *Ruminococcaceae* family members. Finally, only the NCBI database revealed *Barnesiella* as differential, due to the fact that its OTUs were part of the *Unclassified Porphyromonadaceae/uncultured Bacteroidales bacterium/Unclassified S24-7 group* in the other databases.

Contrariwise, taxonomy assignments in the milk microbiota dataset were highly consistent with those reported in the original paper. The four databases equally identified the different *Streptococcus* abundance, as well as the statistically increased *Lactobacillus* and *Pediococcus* in the Rendena breed, *Staphylococcus, Corynebacterium*, and *Unclassified Ruminococcaceae* in the Holstein breed (17). Notably, all but the Greengenes database reported the *Bradyrhizobium* increase in Holstein cows. The *Unclassified Clostridiales* Holstein cows increase was reported by all but SILVA, which identified them as members of the *Christensenellaceae R-7 group, Ruminococcaceae UCG-014* and *UCG-011*. Lastly, only Greengenes found the higher *Unclassified Streptococcaceae* abundance in Rendena cows; the corresponding OTUs were labeled as *Streptococcus, Lactobacillus, Pediococcus, Lactococcus*, and *Unclassified Lactobacillales* in the other databases.

In the extreme longevity dataset, only part of the differentially abundant taxa reported in the original study (18) (*Faecalibacterium, Akkermansia, Eggerthella, Anaerotruncus*) was found by all the databases we used here. Consistent evidence was found for non-significantly altered genera as well: *Bifidobacterium, Bilophila, Odoribacter*, and *Butyricimonas*, the latter two significantly increased in the semi-supercentenarian group in the original paper, but not in our present analysis. Significantly altered taxa (i.e. *Coprococcus, Roseburia, Oscillospira, Christensenellaceae, Synergistaceae*), instead, were only found in the Greengenes database and were missing in the other three, being split into different taxa.

All four databases correctly captured the main differences reported in the original vaginal microbiota paper (19), like the *Lactobacillus* genus depletion in the BV condition, as well as the significant increase of *Gardnerella, Megasphaera, Roseburia, Sneathia, Atopobium*, and *Prevotella*. Relative abundances of these taxa were highly consistent among the four databases and comparable to those of the original paper, with the exception of *Megasphaera* and *Roseburia:* the *Megasphaera* inconsistent relative abundance in the NCBI database, for example, was due to the classification of its OTUs majority as an unclassified member of the *Veillonellaceae* family.

Notably, the original studies considered in the present work were all performed using the 97% Greengenes database, whereas the current comparisons were made using the 100% databases, thus, although we found nearly all the differential genera of the original publications, we could argue that differences were due to the different identity percentages.

### Analysis of taxa misclassification

Some of the taxa inconsistently classified over the four databases were particularly relevant, being the most abundant in the environment (e.g.: *Muribaculum* in mouse infection dataset) or a common finding (e.g. *Roseburia* in human-related microbiota). Thus, we analyzed in detail the discrepancies among these taxa as they could be, potentially, the more problematic to address.

The *Muribaculum* genus constitutes about 30% of the relative abundance of the mouse microbiota. However, it was differently classified in each database: *Unclassified Porphyromonadaceae* (RDP), *Muribaculum* (NCBI), *uncultured bacterium* (*Muribaculaceae* family, SILVA), *Unclassified S24-7* (Greengenes). The *Muribaculaceae* family underwent many major reclassifications from “mouse intestinal bacteria” (27) to an uncultured family of the *Bacteroidales* order and then named after one of the earliest environmental clones belonging to the lineage (S24-7) in both Greengenes and SILVA. The family was lately identified as *Candidatus Homeothermaceae* (28), while *Muribaculum* was not reported in the NCBI repository before July 2016 (29); the *Muribaculaceae* family name has been accepted just recently (30). It is obvious that Greengenes and RDP databases, lastly updated in 2013 and 2016, could not include the latest findings, whereas the *Muribaculaceae* family classification as *Porphyromonadaceae* in RDP was already reported (30). Nonetheless, the SILVA database, which includes the *Muribaculum* genus, classified the majority of these OTUs (14.5% - 29.9% of the total relative abundance) to the upper level of family *Muribaculaceae*, with genus-level *Muribaculum* counting only for 0.5% - 2.8% of that abundance.

In Cremonesi and colleagues’ work (17), the genus *Bradyrhizobium* was found as statistically increased in the Holstein breed when compared to the Rendena. In our present study, this genus was consistently classified by NCBI, RDP, and SILVA; Greengenes reported it as *Nitrobacter*. These two genera, both members of the *Nitrobacteriaceae* family, are closely related, with *Nitrobacter* species often clustered together *Bradyrhizobium japonicum* (31). We observed this inconsistency because Cremonesi and co-workers used the 97% Greengenes, which does not contain *Nitrobacter*, while the Greengenes 100% database used here includes both *Nitrobacter* and *Bradyrhizobium*: our classification, indeed, split the OTUs between *Bradyrhizobium* (for a minor amount only, <0.03% of the average relative abundance) and *Nitrobacter* (3.57% on average).

Both the extreme longevity and the vaginal microbiota datasets revealed an inconsistent classification of the *Roseburia* genus, whose OTUs were classified as either *Roseburia* (Greengenes), *Lachnospiracea incertae sedis* (RDP), *Unclassified Lachnospiraceae* (NCBI), or *Agathobacter* (SILVA). This diversified group accounted for 1.9% and 3.9% of the average relative abundance in the vaginal microbiota and the extreme longevity datasets, respectively, whereas the fraction of the OTUs only classified as *Roseburia* by all databases, was drastically lower (0.2% and 0.7% on average, respectively). The *Roseburia* misclassification can derive from the 16S rRNA gene sequence similarities between with *Agathobacter* (>95%) (32), and the confusion among these two and *Eubacterium*: *Eubacterium rectale* is also classified as *Roseburia rectale* and as *Agathobacter rectalis* (see NCBI:txid39491). The fact that all of these genera belong to the *Lachnospiraceae* family could be pivotal to the issue.

Among incongruently classified taxa in the extreme longevity dataset, we found *Oscillospira*, classified as *Oscillibacter* in NCBI and RDP and as *Ruminococcaceae UCG-002* in SILVA. As a matter of fact, *Oscillibacter* and *Oscillospira* share high similarity in the 16S rRNA sequence (92.6%–92.9% between *Oscillospira guilliermondii* and *Oscillibacter valericigenes*), with the consequence that *Oscillibacter* has often been wrongly assigned to the family *Oscillospiraceae* (persisting up to 2017 in the NCBI taxonomy database (33)). In the SILVA 132 release, oppositely, *Oscillospira* was placed within the *Ruminococcaceae* family and was confounded with the *Ruminococcaceae UGC-002* genus, which encompassed 1816 *uncultured bacteria*. In the latest release (138), however, the genus *Ruminococcaceae UCG-002* was reassigned to the family *Oscillobacteriaceae*.

The concordance among databases in the different taxa classification is related to both the ecosystem complexity and the ecosystem description completeness. The murine microbiota, in fact, was the one with more than half OTUs with discordant taxonomy in all databases, whereas the picture from the human-derived environments was different since the gut microbiota (extreme longevity stool dataset, with its most abundant genera, *Bacteroides*, accounting for <10%) is more diverse than the vaginal one (dominated by the *Lactobacillus* genus); similarly, milk microbiota is dominated by few genera and overall concordance among the four databases is in the order of about 60% of the relative abundance.

### Technical limitations and drawbacks

In this comparison, we preferred to perform all analyses using the same methodology (OTU clustering and QIIME 1.9.0) as in the original work, as we aimed at replicating the exact same procedure with the sole classification database change, without introducing any further confounders.

Nonetheless, we are aware that in the last few years the microbiota analysis pipelines underwent some major modifications and updates: Operational Taxonomic Units (OTUs) now are believed to artificially inflate the number of taxa (34,35) and have been superseded by Amplicon Sequence Variants (ASV). The QIIME pipeline has been discontinued in favor of the more recent QIIME2 (5), and several different alternative analysis pipelines like mothur and UPARSE are currently available. While the best pipeline choice has been analyzed and debated many times (36-39), QIIME is still the most used and well-accepted for microbiome analysis, with more than 22’000 citations.

The last Greengenes update dates back to 2013, whereas no updates have been made on RDP since 2016; SILVA and NCBI are currently the only actively updated. The reduced computational time, together with the fact that it has been the QIIME standard for many years, made Greengenes the most popular and used database. Alternatively, SILVA contains nearly double the number of sequences of Greengenes and resulted very time-consuming for all the four datasets. The direct consequence is that researchers with no access to powerful computational platforms still tend to use Greengenes and miss a decade of taxonomic updates. However, it’s worth noting that using the 97% version SILVA database trimmed down the computational time up to around 10-fold, making this database more feasible and appealing for microbiome projects.

### Comparison with existing literature

Our analysis is the first, to our knowledge, to explore the effects of the database choice on real microbiome samples, representative of different ecosystems and of different sources, from already published papers and publicly available to the scientific community, and the first to assess the eventual alterations in three commonly evaluated aspects of microbiota experiments: the microbiota biodiversity (alpha-diversity), composition (beta-diversity), and composition (relative abundances).

Balvočiūtė and Huson devised an algorithm for mapping the taxonomies from five databases (SILVA, RDP, Greengenes, NCBI, and OTT) one against each other, by mapping them onto a common taxonomy. In general, the number of the intersections was strongly limited by the smallest taxonomy size (Greengenes) whereas SILVA shared the most taxonomic units with NCBI. All other taxonomies mapped well on OTT but, since it has no sequence database associated, its usefulness in the metagenomics context is limited. (4) Park and co-workers used 6 mock samples made up of 59 known equally-distributed bacterial strains DNA and performed taxonomy assignment through UCLUST algorithm against three 16S databases (Greengenes, SILVA, and EzBioCloud), showing that the last one, containing the fewest sequences, performed better than the others regarding richness, evenness, and taxonomic identification (1).

Similarly, Dixit and colleagues employed a collection of 5’895 type-strain, full-length 16S rRNA gene sequences from the RDP website, classified against SILVA, Greengenes, and EzTaxon databases, highlighting discrepancies at various taxonomic levels (3).

Abellan-Schneyder and co-workers performed a nice in-depth comparison, evaluating several variables possibly influencing taxonomic assignments, such as different hypervariable regions, sequence clustering (OTUs *vs*. ASVs), pipeline settings, different databases (Greengenes, RDP, SILVA, GRD, LTP). Comparisons were made on three mock communities, plus DNA extracted from 33 healthy volunteers’ stools. However, the majority of the comparisons were made using the three mock communities and only a few (primer selection and reads truncation effects) using the 33 fecal samples (2).

## CONCLUSION

We here provided evidence about how diversity and taxonomy can be determined by the right database choice. Each database has its advantages and disadvantages, regarding size, computational time, and classification accuracy, and we showed that those factors also depend on the sample dataset of interest, for both the ecosystem complexity and its characterization goodness in the reference databases. Moreover, the usage of recently updated databases is strongly suggested in order to discard some incongruences that may derive from taxonomies based on old or incomplete knowledge. We hope that our comparison study and main taxa reconstruction will help researchers with this crucial choice and improve microbiota investigation.

