## Supplementary Methods, Tables, Figures for "A comparison between Greengenes, SILVA, RDP, and NCBI reference databases in four published microbiota datasets"

Procedures for downloading and formatting the reference databases

#### Ribosomal Database Project (RDP)

Release 11.5 (date) was used (https://rdp.cme.msu.edu/misc/resources.jsp)

Download of the aligned and unaligned sequences

```
wget https://rdp.cme.msu.edu/download/current_Bacteria_unaligned.fa.gz
--no-check-certificate
wget https://rdp.cme.msu.edu/download/current_Bacteria_aligned.fa.gz --no-
check-certificate
```

Keep only the interesting fields (ID and sequence)

```
gzip -dc current_Bacteria_unaligned.fa.gz| awk '{print $1}'
>current_Bacteria_RDP_11_5.fna
gzip -dc current_Bacteria_aligned.fa.gz| awk '{print $1}'
>current_Bacteria_RDP_11_5.aligned.fna
```

Format of the taxonomy in order to be used by QIIME

```
grep ">" current_Bacteria_unaligned.fa >temp_taxa

sed 's/, member of the beta subclass//' temp_taxa|sed
's/Lineage=Root;rootrank;//;s/Bacteria;domain;//;s/\"//g'|sed
's/phylum;/phylum\t/;s/class;/class\t/;s/subclass;/subclass\t/;s/order;/ord
er\t/;s/suborder;/suborder\t/;s/family;/family\t/'|awk
'BEGIN{FS=OFS="\t";ORS=""}{for(i=1;i<=NF;i++){if($i !~ /subclass/ && $i !
~ /suborder/) print $i"\t"};print "\n"}'|perl -naF"\t" -e '($ph=$1) if ($_
=~ /\t(\w+)\;phylum/); ($cl=$1) if ($_ =~ /\t(\w+)\;class/); ($or=$1) if
($_ =~ /\t(\w+)\;order/); ($fam=$1) if ($_ =~ /\t(\w+)\;family/);($gen=$1)
if ($_ =~ /\t(\w+)\;genus/); ($name=$1 and $sp=$2 and $str=$3) if ($F[0] =~
/^>(\w+)\s?(.*)/); print
"$name\tk__Bacteria;p__$ph;c__$cl;o__$or;f__$fam;g__$gen;s__$sp;\n";$ph="";
$cl="";$or="";$fam="";$gen="";$sp="";$name=""'|perl -pae '$_ =~ s/;\s.+;
$//'|sed 's/;$//;s/\*//' >RDP_11_5.id2taxa.txt
```

Lanemask file to be used for filtering out the positions composed by all gaps (meaningless) in multiple alignments

```
find_all_gap_positions.py current_Bacteria_RDP_11_5.aligned.fa
lanemask_in_1s_and_0s
```

#### National Center for Biotechnology Information (NCBI)

Release 17/06/2018 was used (ftp://ftp.ncbi.nlm.nih.gov/blast/db/). At present, the database has been updated on 01/03/2022 and includes 22311 sequences

In order to associate sequence IDs to taxonomies, the utilities in Entrez_qiime GitHub project (https://bakerccm.github.io/entrez_qiime/) have been downloaded

```
wget https://github.com/bakerccm/entrez_qiime/archive/v2.0.tar.gz
tar -xf v2.0.tar.gz
```

Download of the 16S reference database from NCBI FTP site

```
wget ftp://ftp.ncbi.nlm.nih.gov/blast/db/16SMicrobial.tar.gz
tar -xf 16SMicrobial.tar.gz
```

Starting from the Blast-formatted database, rebuild the FASTA file by using the blastdbcmd from BLAST+

```
blastdbcmd -entry all -db 16SMicrobial -out /dev/stdout|awk '{print $1}' >
16SMicrobial.fna
```

Following the instructions in https://github.com/bakerccm/entrez_qiime/blob/master/entrez_qiime.pdf, download the files needed to rebuild the sequence taxonomies

```
wget ftp://ftp.ncbi.nlm.nih.gov/pub/taxonomy/taxdump.tar.gz
wget
ftp://ftp.ncbi.nlm.nih.gov/pub/taxonomy/accession2taxid/nucl_gb.accession2t
axid.gz

gunzip nucl_gb.accession2taxid.gz &
tar -xf taxdump.tar.gz
```

Use of entrez_qiime.py utility to generate the taxonomy compatible with QIIME pipeline

```
python2.7 entrez_qiime-2.0/entrez_qiime.py -i 16SMicrobial.fna -o
./temp.txt -g ./log_entrez_qiime.log -n . -a nucl_gb.accession2taxid

# Add the field “Bacteria” as the kingdom level taxonomy
sed 's/ /_/g' temp.txt|awk '{print $1"\tBacteria;"$2}'
>16SMicrobial.id2taxa.txt

rm temp.txt
```

Generate the multiply-aligned sequence file by using ClustalOmega tool (http://www.clustal.org/omega/#Download)

```
clustalo -i 16SMicrobial.fna -t DNA --threads=40 -o
16SMicrobial.aligned.fna --log=log_clustalo.log -v --outfmt=fasta --force
&>errs
```

Lanemask file to be used for filtering out the positions composed by all gaps (meaningless) in multiple alignments

```
find_all_gap_positions.py 16SMicrobial.aligned.fna lanemask_in_1s_and_0s
```

#### Greengenes

Releases 13_5 (raw data) and 13_8 (97% clustered data) were used (ftp://greengenes.microbio.me/greengenes_release/). Release 13_8 (last

Greengenes update) contains an udate of 13_5 in terms of OTUs and clustering. Thus, raw FASTA sequences of the database are those of 13_5 release

##### Unclustered raw data (100% identity)

```
wget
ftp://greengenes.microbio.me/greengenes_release/gg_13_5/gg_13_5.fasta.gz
wget
ftp://greengenes.microbio.me/greengenes_release/gg_13_5/gg_13_5_taxonomy.tx
t.gz
wget
ftp://greengenes.microbio.me/greengenes_release/gg_13_5/gg_13_5_ssualign.fa
sta.gz
```

Lanemask file to be used for filtering out the positions composed by all gaps (meaningless) in multiple alignments

```
find_all_gap_positions.py gg_13_5_ssualign.fasta lanemask_in_1s_and_0s
```

##### 97% clustered data

```
ftp://greengenes.microbio.me/greengenes_release/gg_13_8_otus/taxonomy/97_ot
u_taxonomy.txt
ftp://greengenes.microbio.me/greengenes_release/gg_13_8_otus/rep_set_aligne
d/97_otus.fasta
ftp://greengenes.microbio.me/greengenes_release/gg_13_8_otus/rep_set/97_otu
s.fasta
```

Lanemask file is the one provided within the QIIME 1.9.0 package

#### ARB-SILVA ribosomal RNA database project (SILVA)

Release 132 was used (https://www.arb-silva.de/no_cache/download/archive/qiime/). This is the last SILVA release with provided pre-formatted files for QIIME.

Download of QIIME-formatted SILVA 132 database. This archive contains both the raw sequences and the clustered databases and taxonomies

```
wget https://www.arb-
silva.de/fileadmin/silva_databases/qiime/Silva_132_release.zip
unzip Silva_132_release.zip
```

Download of the multiple alignment of the full database

```
wget https://www.arb-
silva.de/fileadmin/silva_databases/release_132/Exports/SILVA_132_SSURef_tax
_silva_full_align_trunc.fasta.gz
gunzip SILVA_132_SSURef_tax_silva_full_align_trunc.fasta.gz
```

##### 97% clustered data

For the 97% clustered data, format the taxonomy in order to be compatible with QIIME, filtering out ambiguous data and substituting the phylogenetic level indication “D_0” -> “k__” and so on.

```
{
grep -v "Ambiguous_taxa;D_"
taxonomy/16S_only/97/majority_taxonomy_7_levels.txt
grep "Ambiguous_taxa;D_6"
taxonomy/16S_only/97/majority_taxonomy_7_levels.txt
}|sed 's/Ambiguous_taxa;D_6.*/Ambiguous_taxa;Ambiguous_taxa/'|sed
's/D_0/k/;s/D_1/p/;s/D_2/c/;s/D_3/o/;s/D_4/f/;s/D_5/g/;s/D_6/s/'|perl -nae
'$count=0;($count++) while ($_ =~ m/Ambiguous/g);print "$count\t$_";'|awk
'BEGIN{FS=OFS="\t"}{if($1==1){sub(/Ambiguous_taxa.*/,"s__",$3)}else if
($1==2){sub(/Ambiguous_taxa;Ambiguous_taxa.*/,"g__;s__",$3)} else if
($1==3){sub(/Ambiguous_taxa;Ambiguous_taxa;Ambiguous_taxa.*/,"f__;g__;s__",
$3)} else if ($1==4)
{sub(/Ambiguous_taxa;Ambiguous_taxa;Ambiguous_taxa;Ambiguous_taxa.*/,"o__;f
__;g__;s__",$3)} else if ($1==5)
{sub(/Ambiguous_taxa;Ambiguous_taxa;Ambiguous_taxa;Ambiguous_taxa;Ambiguous
_taxa.*/,"c__;o__;f__;g__;s__",$3)} else if ($1==6)
{sub(/Ambiguous_taxa;Ambiguous_taxa;Ambiguous_taxa;Ambiguous_taxa;Ambiguous
_taxa;Ambiguous_taxa.*/,"p__;c__;o__;f__;g__;s__",$3)} else if ($1==7)
{sub(/"Ambiguous_taxa;Ambiguous_taxa;Ambiguous_taxa;Ambiguous_taxa;Ambiguou
s_taxa;Ambiguous_taxa;Ambiguous_taxa.*/,"k__;p__;c__;o__;f__;g__;s__",$3)};
print $0}'|cut -f2- >taxonomy/16S_only/97/97_otu_taxonomy_QIIME.txt
```

Lanemask file to be used for filtering out the positions composed by all gaps (meaningless) in multiple alignments

```
find_all_gap_positions.py rep_set_aligned/97/97_alignment.fna
rep_set_aligned/97/lanemask_in_1s_and_0s
```

##### Unclustered raw data (100% identity)

In order to properly format the raw (unclustered) data, download the necessary code from Mike Robson’s Github (https://github.com/mikerobeson/Misc_Code/tree/master/SILVA_to_RDP)

```
wget
https://raw.githubusercontent.com/mikerobeson/Misc_Code/master/SILVA_to_RDP
/prep_silva_data.py
wget
https://raw.githubusercontent.com/mikerobeson/Misc_Code/master/SILVA_to_RDP
/prep_silva_taxonomy_file.py
wget
https://gist.githubusercontent.com/walterst/0a4d36dbb20c54eeb952/raw/7a960f
4e0550bde24896e5c3eee2d31ca380fdf1/parse_nonstandard_chars.py
wget
https://gist.githubusercontent.com/walterst/9ddb926fece4b7c0e12c/raw/487170
e6a02752eacf894fd9d1aace6832b55968/parse_to_7_taxa_levels.py
```

**Reference sequences**

Deletion of non-standard characters from SILVA reads (gaps-related (".","-"), spaces (" ")) and converts “U" to "T"

```
fasta2tabular.sh initial_reads_SILVA132.fna |awk 'BEGIN{FS=OFS="\t"}
{gsub("U","T",$2);gsub("\.","",$2);gsub("-","",$2);gsub(" ","",$2); print
">"$1"\n"$2}' >t1.fna
```

Starting from the sequence file, generates the final FASTA file and extracts the taxonomies

```
python prep_silva_data.py t1.fna taxa_out.txt
initial_reads_SILVA132.QIIME.fna
```

**Taxonomies**

Pruning of characters which give errors in RDP classifier (such as "*") from the taxonomy file

```
python parse_nonstandard_chars.py taxa_out.txt > taxa_out.clean.txt
```

Formatting of the file in order to be compatible with RDP classifier

```
python prep_silva_taxonomy_file.py taxa_out.clean.txt
taxa_out.clean.RDP.txt
```

Extracting a 7-level taxonomy

```
python parse_to_7_taxa_levels.py taxa_out.clean.RDP.txt
taxa_out.clean.RDP.7levels.txt
```

Re-format of taxonomy. Delete all classification beyond level 6 (species) and substitute the phylogenetic level indication “D_0” -> “k__” and so on.

```
sed 's/D_0/k/;s/D_1/p/;s/D_2/c/;s/D_3/o/;s/D_4/f/;s/D_5/g/;s/D_6/s/'
taxa_out.clean.RDP.7levels.txt >raw_taxonomy_QIIME.txt
```

Due to the high number of sequences within the raw database file, the use of lanemask file is not possible (computationally too intensive). Thus, the procedure for filtering out positions due to entropy and gap fractions. Recommended parameters are those suggested in the README file of the SILVA132 release for QIIME (i.e.: -e 0.10, -g 0.80)

**Multi-aligned sequences file**

Re-format of multiply-aligned FASTA file. Convert “U” to “T” and “.” to “-”

```
fasta2tabular.sh /storage-
daredevil/qiime_resources/SILVA_132_QIIME_release/raw_data/SILVA_132_SSURef
_tax_silva_full_align_trunc.fasta |awk 'BEGIN{FS=OFS="\t"}{gsub("U","T",
$2);gsub("\.","-",$2);split($1,a," ");print ">"a[1]"\n"$2}' >/storage-
daredevil/qiime_resources/SILVA_132_QIIME_release/raw_data/SILVA_132_SSURef
_tax_silva_full_align_trunc.perQIIME.fasta
```

### SUPPLEMENTARY FIGURES

#### Supplementary Figure 1.

Rarefaction curves of alpha-diversity estimations according to Faith's phylogenetic diversity metric (PD whole tree). For each of the four datasets, curves obtained using SILVA and Greengenes (GG) databases at 97% or 100% identity are shown. Lines indicate average alpha-diversity with standard deviation-based error bars.

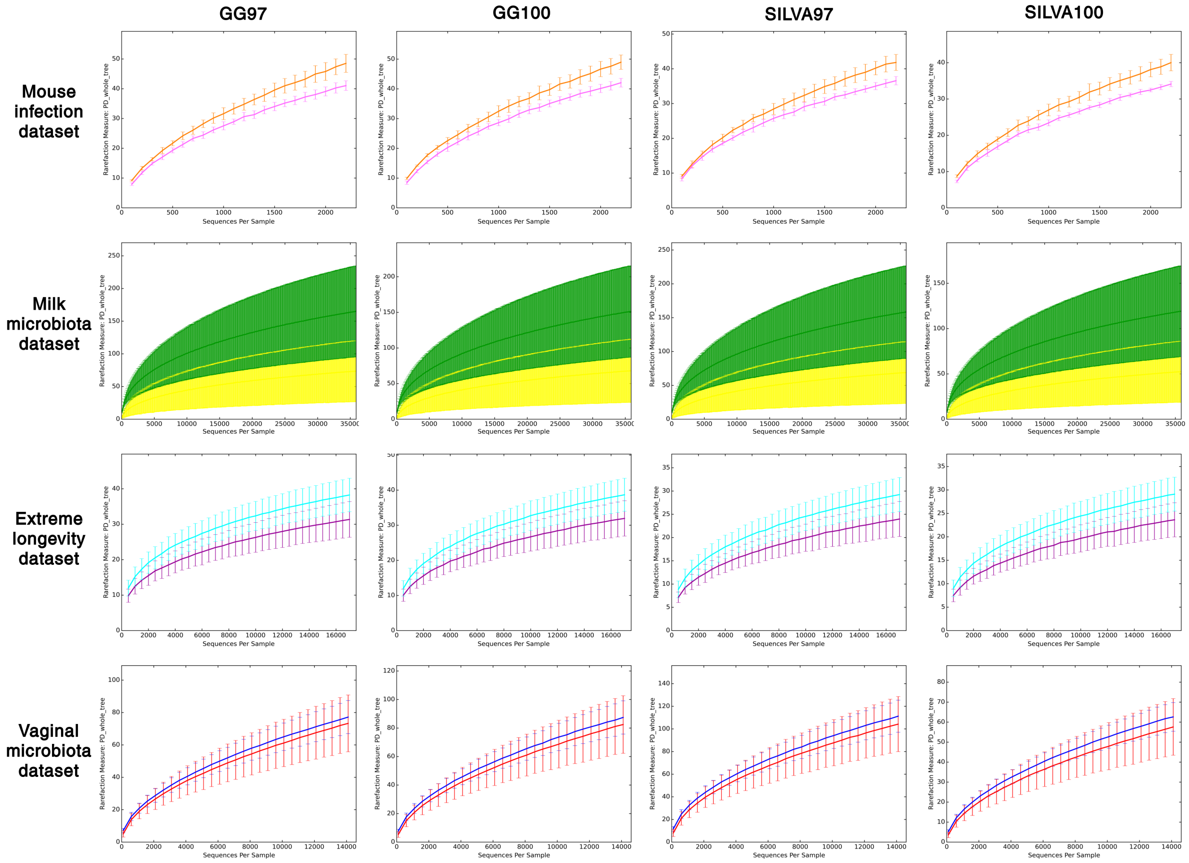

#### Supplementary Figure 2.

(**A**) Barplots representing the relative abundance at genus level whose classification was concordant among the four databases over the four datasets; (**B**) Same barplots as in (A), but comprising only genera completely classified at genus level (i.e.: no database reporting a “unclassified” taxonomy); (**C**) Same barplots as in (A), but comprising only genera with at least one database reporting an “unclassified” taxonomy at genus level.

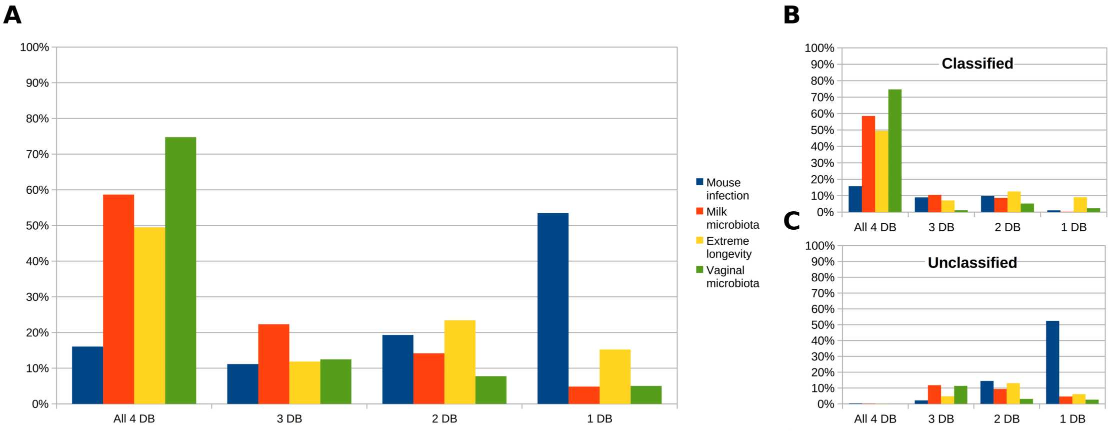

### SUPPLEMENTARY TABLES

#### Supplementary Table 1.

Comparison between PD whole tree alpha diversity metric between SILVA and Greengenes. For each dataset, database and percentage of identity, the average value ± standard deviation, together with the p-value from a Mann-Whitney U-test is reported. * indicate statistical significance (p<0.05)

| Dataset | n | % identity | SILVA | Greengenes | p-value |
|---|---|---|---|---|---|
| Extreme longevity | 39 | 97 | 27.2 ± 4.5 | 35.6 ± 5.9 | <0.0001* |
| | | 100 | 27.0 ± 4.5 | 36.1 ± 5.8 | <0.0001* |
| Milk microbiota | 30 | 97 | 257.0 ± 149.6 | 268.9 ± 152.7 | 0.7615 |
| | | 100 | 193.8 ± 110.4 | 246.8 ± 140.6 | 0.1098 |
| Mouse infection | 12 | 97 | 78.4 ± 6.4 | 89.6 ± 8.9 | 0.0018* |
| | | 100 | 74.1 ± 6.8 | 91.1 ± 8.0 | <0.0001* |
| Vaginal microbiota | 41 | 97 | 107.7 ± 20.3 | 75.2 ± 14.6 | <0.0001* |
| | | 100 | 60.0 ± 11.6 | 84.9 ± 16.8 | 0.0001* |

#### Supplementary Table 2.

Comparison of Unifrac distances between databases at different percentages of identity. For both Greengenes and SILVA, the average of unweighted (top) or weighted (bottom) Unifrac distances ± standard deviation is reported, together with the p-value from a two-tailed unpaired t-test. * indicate statistical significance (p<0.05)

| Unweighted Unifrac distances | | | | | | |
|---|---|---|---|---|---|---|
| | Greengenes | | | SILVA | | |
| | **100%** | **97%** | **p-value[a]** | **100%** | **97%** | **p-value** |
| Extreme longevity | 0.69±0.04 | 0.67±0.04 | <0.001* | 0.69±0.04 | 0.69±0.04 | 0.648 |
| Milk microbiota | 0.75±0.07 | 0.75±0.07 | 0.608 | 0.74±0.06 | 0.75±0.07 | 0.379 |
| Mouse infection | 0.66±0.05 | 0.66±0.05 | 0.619 | 0.63±0.05 | 0.66±0.05 | 0.084 |
| Vaginal microbiota | 0.75±0.05 | 0.75±0.05 | 0.224 | 0.76±0.05 | 0.73±0.05 | <0.001* |

| Weighted Unifrac distances | | | | | | |
|---|---|---|---|---|---|---|
| | Greengenes | | | SILVA | | |
| | **100%** | **97%** | **p-value** | **100%** | **97%** | **p-value** |
| Extreme | 0.31±0.0 | 0.29±0. | <0.001 | 0.21±0. | 0.20±0. | 0.001* |

| | | | | | | |
|---|---|---|---|---|---|---|
| longevity | 7 | 06 | * | 05 | 05 | |
| Milk microbiota | 0.24±0.11 | 0.28±0.13 | <0.001 * | 0.20±0.09 | 0.25±0.11 | <0.001 * |
| Mouse infection | 0.20±0.06 | 0.20±0.07 | 0.945 | 0.17±0.05 | 0.30±0.09 | <0.001 * |
| Vaginal microbiota | 0.35±0.18 | 0.31±0.18 | <0.001 * | 0.22±0.13 | 0.52±0.24 | <0.001 * |

[a] p-value from unpaired two-tailed t-test

#### Supplementary Table 3.

Comparison of Unifrac distances among the four taxonomy databases. For all databases, the average of unweighted (top) or weighted (bottom) Unifrac distances ± standard deviation is reported, together with the p-value from a one-way ANOVA test, followed by a Tukey HSD test for pairwise comparisons. * indicate statistical significance (p<0.05).

| Unweighted Unifrac distances | | | | | | |
|---|---|---|---|---|---|---|
| | **Greengenes** | **NCBI** | **RDP** | **SILVA** | **p-value [a]** | **Pairwise tests [b]** |
| Extreme longevity | 0.69±0.04 | 0.67±0.05 | 0.67±0.04 | 0.69±0.04 | <0.001 * | 1-2,1-3, 2-4, 3-4 |
| Milk microbiota | 0.75±0.07 | 0.74±0.07 | 0.75±0.07 | 0.74±0.06 | 0.288 | -- |
| Mouse infection | 0.66±0.05 | 0.67±0.05 | 0.63±0.05 | 0.63±0.05 | 0.026 | -- |
| Vaginal microbiota | 0.75±0.05 | 0.73±0.05 | 0.73±0.05 | 0.76±0.05 | <0.001 * | 1-2,1-3, 2-4, 3-4 |

| Weighted Unifrac distances | | | | | | |
|---|---|---|---|---|---|---|
| | **Greengenes** | **NCBI** | **RDP** | **SILVA** | **p-value** | **Pairwise tests** |
| Extreme longevity | 0.31±0.07 | 0.42±0.12 | 0.29±0.07 | 0.21±0.05 | <0.001 * | All |
| Milk microbiota | 0.24±0.11 | 0.41±0.18 | 0.25±0.11 | 0.20±0.09 | <0.001 * | All but 1-3 |
| Mouse infection | 0.20±0.06 | 0.25±0.08 | 0.31±0.09 | 0.17±0.05 | <0.001 * | 1-3, 2-3, 2-4, 3-4 |
| Vaginal microbiota | 0.35±0.18 | 0.57±0.27 | 0.50±0.26 | 0.22±0.13 | <0.001 * | All |

[a] p-value from one-way ANOVA test
[b] performed by Tukey HSD test
